# Selection and validation of reference genes for quantitative Real-Time PCR in *Arabis alpina*

**DOI:** 10.1101/517367

**Authors:** Lisa Stephan, Vicky Tilmes, Martin Hülskamp

**Affiliations:** Botanical Institute, Biocenter, Cologne University, Zülpicher Straße 47b, 50674 Cologne, Germany; Department of Plant Developmental Biology, Max Planck Institute for Plant Breeding Research, Carl von Linne Weg 10, 50829 Cologne, Germany

## Abstract

*Arabis alpina* is a perennial arctic-alpine plant and an upcoming model organism for genetics and molecular biology for the Brassicaceae family. One essential method for most molecular approaches is the analysis of gene expression by quantitative Real-Time PCR (qPCR). For the normalisation of expression data in qPCR experiments, it is essential to use reliable reference genes that are not affected under a wide range of conditions. In this study we establish a set of 15 *A. alpina* reference genes that were tested under different conditions including cold, drought, heat, salt and gibberellic acid treatments. Data analyses with geNORM, BestKeeper and NormFinder revealed the most stable reference genes for the tested conditions: *RAN3, HCF* and *PSB33* are most suitable for cold treatments; *UBQ10* and *TUA5* for drought; *RAN3, PSB33* and *EIF4a* for heat; *CAC, TUA5, ACTIN 2* and *PSB33* for salt and *PSB33* and *TUA5* for gibberellic acid treatments. *CAC* and *ACTIN 2* showed the least variation over all tested samples. In addition, we show that two reference genes are sufficient to normalize qPCR data under our treatment conditions. In future studies, these reference genes can be used for an adequate normalisation and thus help to generate high quality qPCR data in *A. alpina*.

## Introduction

During the last years, *A. alpina* has been established as a new model system in the Brassicaceae family (1,2). It is native to mountains and arctic-alpine habitats (3,4) and combines several features enabling genetic and molecular studies: it is diploid, self-fertile, has a small and sequenced genome and can be transformed with *Agrobacterium tumefaciens* (1). *A. alpina* has an evolutionary distance to *A. thaliana* of about 26 to 40 million years (4,5). This facilitates functional comparisons of biological processes, as orthologous genes can be identified by sequence similarity and synteny (6).

Most molecular studies require quantitative analyses of the expression of genes of interest by quantitative Real-Time PCR. For proper comparisons of expression levels, the expression data of the genes under study are normalized using genes as a reference that show no or very little variation under different conditions. In 2009, the Minimum Information for Publication of Quantitative Real-Time PCR Experiments (MIQE) guidelines were published, with the aim to provide a consensus on correct performance and interpretation of qPCR experiments (Bustin et al., 2009). These guidelines should ensure that the normalisation enables the comparison of transcripts in different samples by correcting variations in yields of extraction and reverse transcription and the efficiency of amplification. A pre-requisite for any qPCR analysis are suitable primer sets for reference genes that are thoroughly tested. These need to fulfil various requirements: primers should create a specific amplicon of 80 to 200 bp, without creating primer dimers. The amplification should be carried out with close to 100 % efficiency and show a linear standard curve with a correlation of more than 0.99. In general, there should be minimal variation between replicates, indicating consistent performance of the primers.

In this study we established primer pairs for 15 reference genes that can be used for future qPCR studies in *A. alpina*: *ADENOSINE TRIPHOSPHATASE* (*ATPase*), *THIOREDOXIN, HIGH CHLOROPHYLL FLUORESCENCE 164* (*HCF*), *EUKARYOTIC TRANSLATION INITIATION FACTOR 4A1* (*EIF4a*), *RAN GTPASE 3* (*RAN3*), *UBIQUITIN 10* (*UBQ10*), *ACTIN 2, PHOTOSYSTEM B PROTEIN 33* (*PSB33*), *HISTONE H3, NAD(P)H PLASTOQUINONE DEHYDROGENASE COMPLEX SUBUNIT O* (*NdhO*), *TUBULIN ALPHA 5* (*TUA5*), *18s RIBOSOMAL RNA* (*18srRNA*), *CLATHRIN ADAPTOR COMPLEX MEDIUM SUBUNIT* (*CAC*), *SAND family protein* (*SAND*) and *HEAT SHOCK PROTEIN 81.2/90* (*HSP81.2/90*). The primers were thoroughly tested, and reference genes were evaluated for variations in their expression under different conditions including cold, drought, heat and salt in whole seedlings and gibberellic acid (GA) treatments in leaves. Using genes specifically responding to the different treatment, we demonstrate the impact of normalization with our reference genes.

## Results

### Selection and validation of reference genes

To compile a set of suitable reference genes, we pursued three approaches: first, we selected *A. thaliana* genes that are known to show little variation under different conditions. Corresponding orthologs in *A. alpina* were then identified by sequence similarity and synteny. Second, we identified genes which show stable expression in *A. alpina* over an extended period of time. Third, we included a well-established reference gene from *A. alpina* from former studies. Thus, we created a set of 15 reference genes (Table 1), including nine orthologs to *A. thaliana* reference genes, five novel reference genes and *RAN3*, a known reference gene for qPCR in *A. alpina* (1). In a first step, we amplified the gene fragments and verified the amplicon by sequencing. In addition, we analysed the melting curves to exclude unspecific products and/or primer dimers (Fig S1).

**Table 1.**
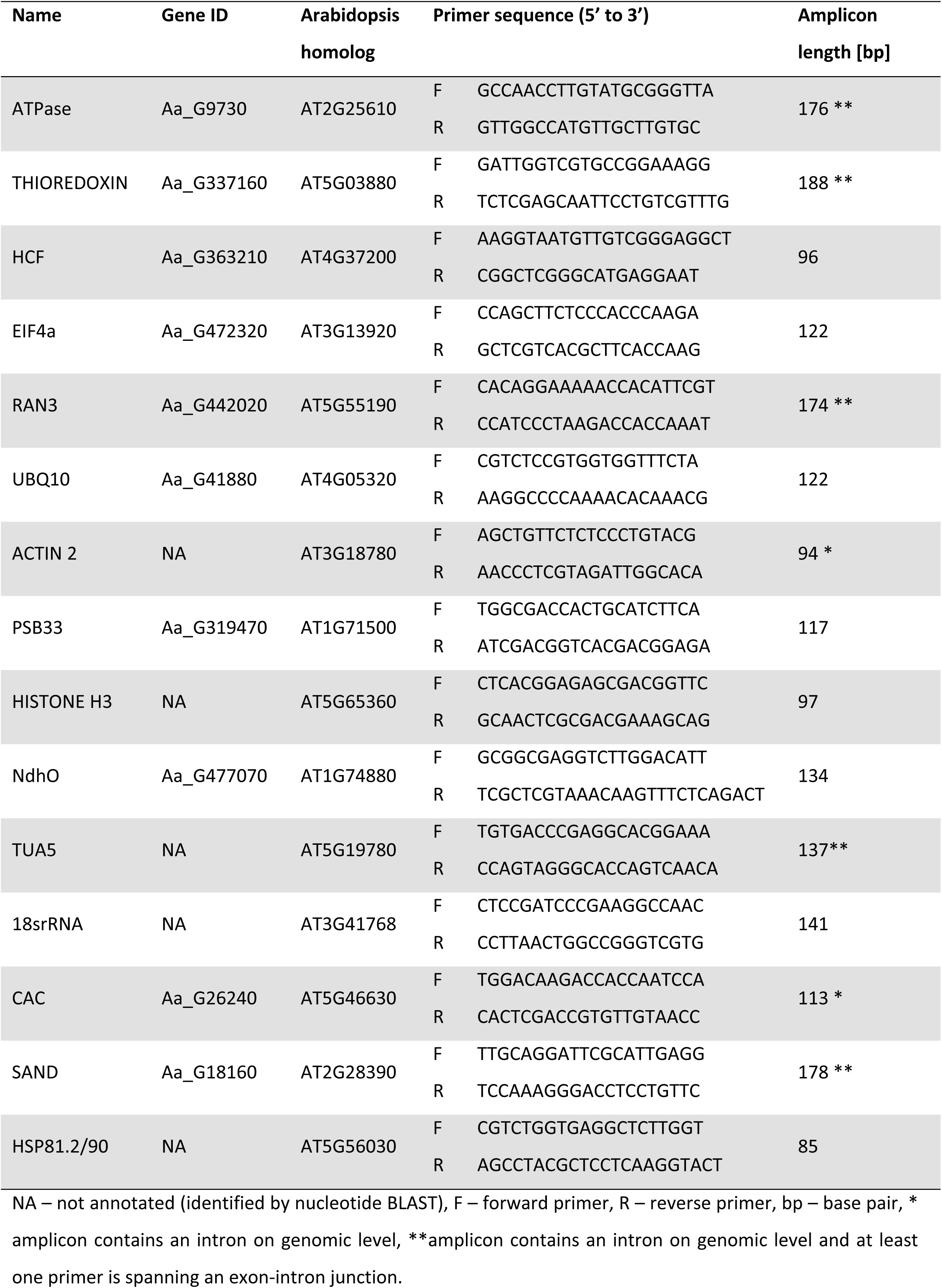
**Candidate reference genes, primers and amplicons.**

Second, we determined the primer efficiencies and correlation coefficients to demonstrate the quality of the primer pairs (Table 2, Fig S2). All reference gene primers displayed an efficiency between 96.42 and 107.01 % and a correlation higher than 0.99.

**Table 2.** **Primer efficiencies and correlation of reference gene candidates.**

| Gene | Efficiency [%] | R<sup>2</sup> |
| --- | --- | --- |
| ATPase | 98.33 | -0.996 |
| THIOREDOXIN | 101.92 | -0.992 |
| HCF | 99.98 | -0.997 |
| EIF4a | 107.01 | -0.992 |
| RAN3 | 102.30 | -0.998 |
| UBQ10 | 102.52 | -0.999 |
| ACTIN 2 | 102.85 | -0.999 |
| PSB33 | 99.51 | -1.000 |
| HISTONE H3 | 99.14 | -0.999 |
| NdhO | 94.34 | -0.999 |
| TUA5 | 98.32 | -1.000 |
| 18srRNA | 98.65 | -1.000 |
| CAC | 98.07 | -0.999 |
| SAND | 96.42 | -0.999 |
| HSP81.2/90 | 96.74 | -0.999 |

Efficiency [%] – efficiency of the primers was calculated with $E[\%] = 100 \times (10^{\left(-\frac{1}{slope}\right)} - 1)$ , slope – average Cq values (y) and log10 values of six serial dilutions (x) were used to calculate the slope of a regression line with the formula $slope = \frac{\sum(x - \bar{x})(y - \bar{y})}{\sum(x - \bar{x})^2}$ , R<sup>2</sup> – correlation of the Cq values calculated with the formula $\rho_{x,y} = \frac{Cov(X,Y)}{\sigma_x \times \sigma_y}$ .

### Expression stability of reference genes after treatments

The expression levels of the reference genes were detected as cycle quantification (Cq) values. The mean Cq values over all treatments ranged between 15.10 (*18srRNA*) and 26.68 (*SAND*). The *18srRNA, UBQ10* and *ACTIN 2* genes showed the highest expression/lowest Cq (Fig 1). Overall, Cq values for a single reference gene varied 4.30 on average, with a minimum variation of 2.82 (*UBQ10*) and a maximum variation of 8.12 (*HSP81.2/90*).

**Fig 1.**
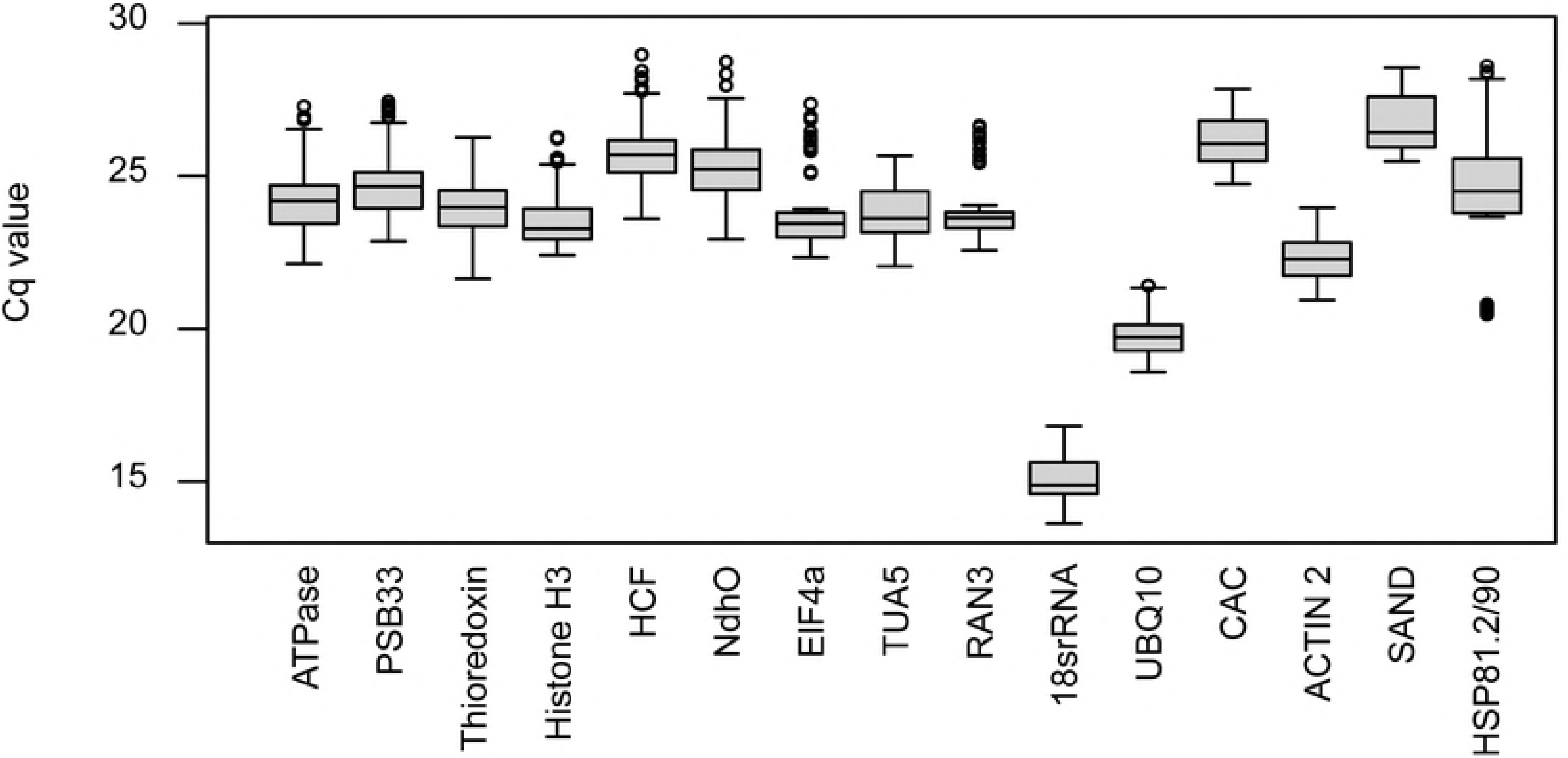
qPCR Cq values of candidate reference genes in all treatments. The box includes all data points between the 25 % quantile and the 75 % quantile. The whiskers span all values below and above, excluding outliers which are displayed as dots. The median, which marks the 50 % quantile, is displayed as a line.

To determine to what extend the selected reference genes respond to different stimuli, we analysed their expression in cold, drought, heat, salt and gibberellic acid treatments. The stability was calculated using three common statistical algorithms: NormFinder (7), BestKeeper (8) and geNorm (9). These algorithms can be used to rank reference genes of a given set by their stability and determine the most stable genes for the tested conditions. In addition, geNorm provides a cut off value of 1.5, above which primer pairs are considered adequate for normalisation. All genes in all treatments, except for *HSP81.2/90* under heat stress conditions, met this criterion. Thus, all reference genes reported here can be considered suitable for a wide range of conditions. The most stable genes for cold treatments were *RAN3, HCF* and *PSB33*. According to the BestKeeper algorithm, *EIF4a* could be considered as a reference for cold treatments as well. However, especially geNorm advises against this gene, ranking it at position eight. After drought treatment, *UBQ10* and *TUA5* showed the least overall variation in the three methods. Here in particular, the results strongly differed between the algorithms. While *HISTONE H3* ranked first for BestKeeper, it was found to be least stable in the other two methods. On the contrary, *EIF4a* and *THIOREDOXIN* showed high stability in NormFinder and geNorm, but low values in BestKeeper. Under heat conditions, *RAN3, PSB33* and *EIF4a* were most stable over all algorithms and under salt conditions *CAC, TUA5, ACTIN 2* and *PSB33* performed best. Finally, the GA treatments caused least changes of the expression of *PSB33* and *TUA5*, while *THIOREDOXIN* showed stable expression only for NormFinder and geNorm. When taking all stress treatments into account, *THIOREDOXIN* and *CAC* showed the least variation. When considering all treatments, *CAC* and *ACTIN 2* showed the most stable expression (Table S2).

### Optimal number of reference genes

For the optimal normalisation, it is necessary to use two or more reference genes in each experiment. The optimal number of reference genes can be determined with the geNorm algorithm, which calculates the pairwise variation V_n/n+1_ based on the normalisation factors NF_n_ and NF_n+1_, with n≥2. If V_n/n+1_ is below 0.15, n is the optimal number of reference genes. For all tested treatments, individually or combined, two reference genes are sufficient to normalise qPCR measurements (Fig 2). The use of a third reference would not improve the results significantly.

**Fig 2.**
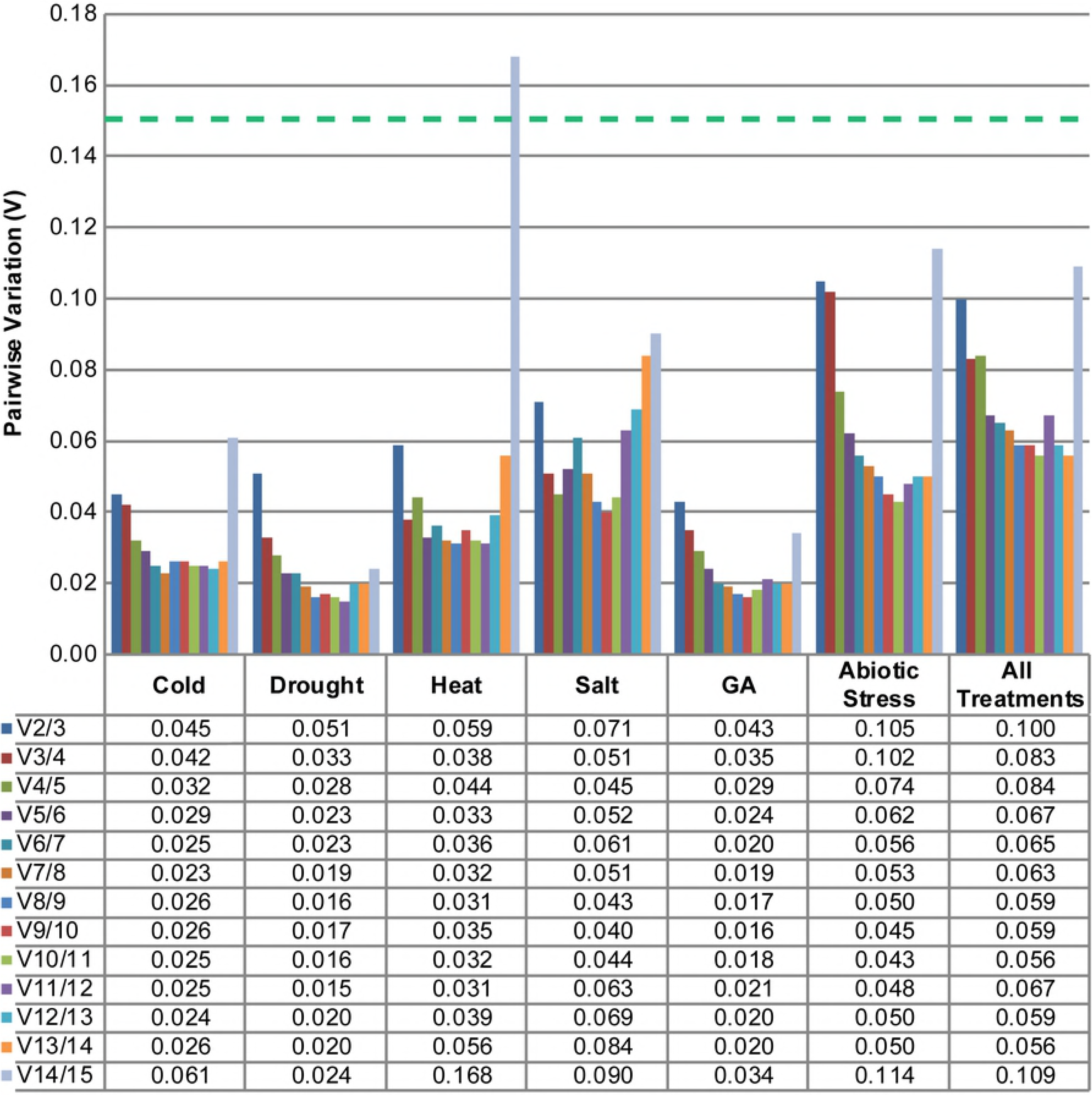
Optimal number of reference genes for various conditions. The geNorm algorithm was used to determine the pairwise variation (V) between the reference genes for treatments with cold, drought, heat, salt and gibberellic acid. The threshold for adequate normalisation is V≤0.15, indicated by the green dashed line.

### Impact of reference genes on the normalisation of samples in abiotic stress and hormone treatments

The efficiencies of treatments with cold, drought, heat, salt and gibberellic acid were controlled by the expression analysis of specific stress response genes by qPCR using primers for *RD29A* (*RESPONSIVE TO DESICCATION 29A*, cold and drought responsive), *HSP81.2/90* (heat responsive), *TSPO* (*OUTER MEMBRANE TRYPTOPHAN-RICH SENSORY PROTEIN-RELATED*, salt responsive) and *GA3ox1* (*GIBBERELLIN 3-OXIDASE 1*, GA responsive). All primers showed efficiencies of 80.28 to 104.22 % and correlations of more than 0.99 (Table 4).

**Table 3.**
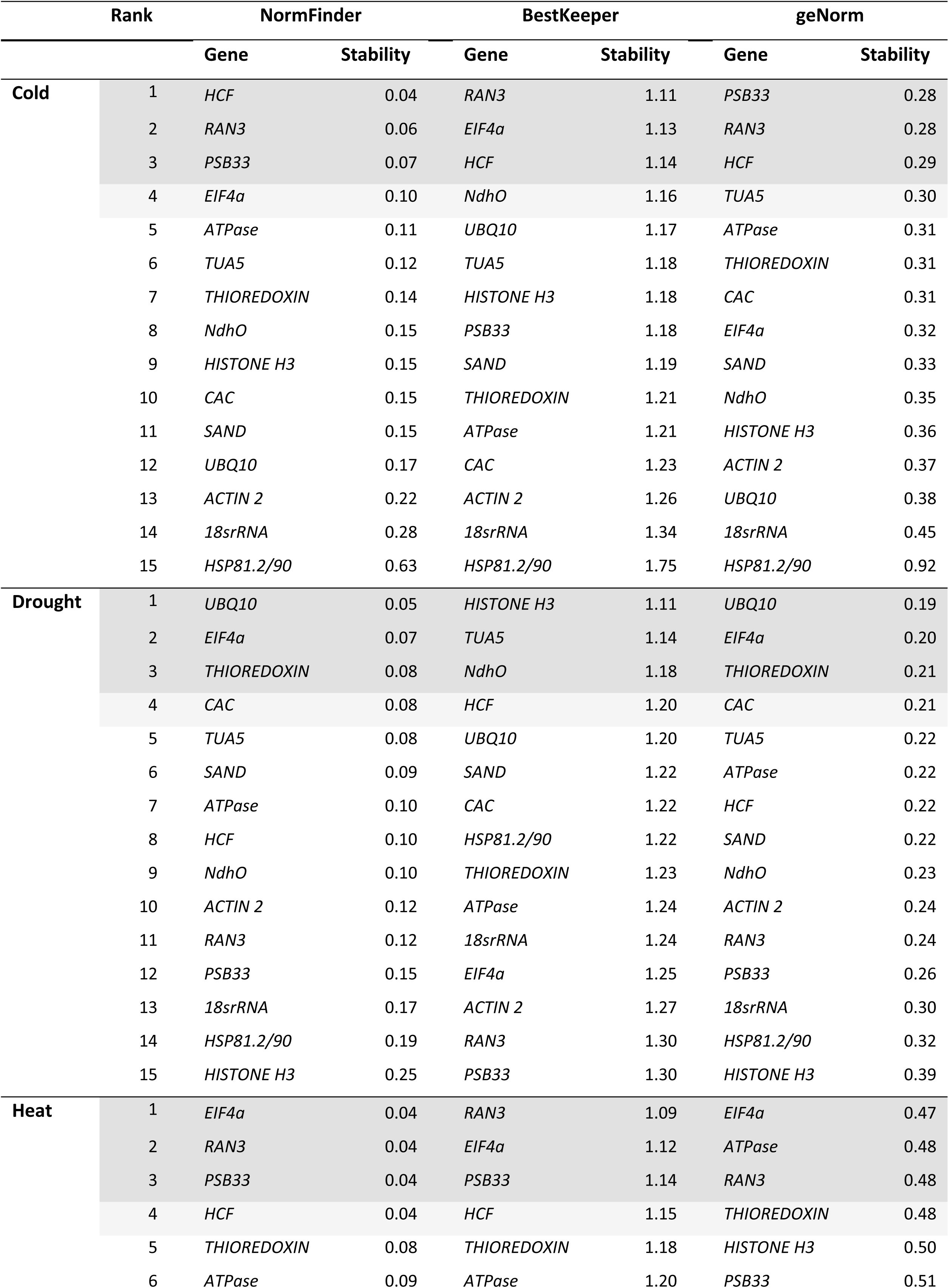
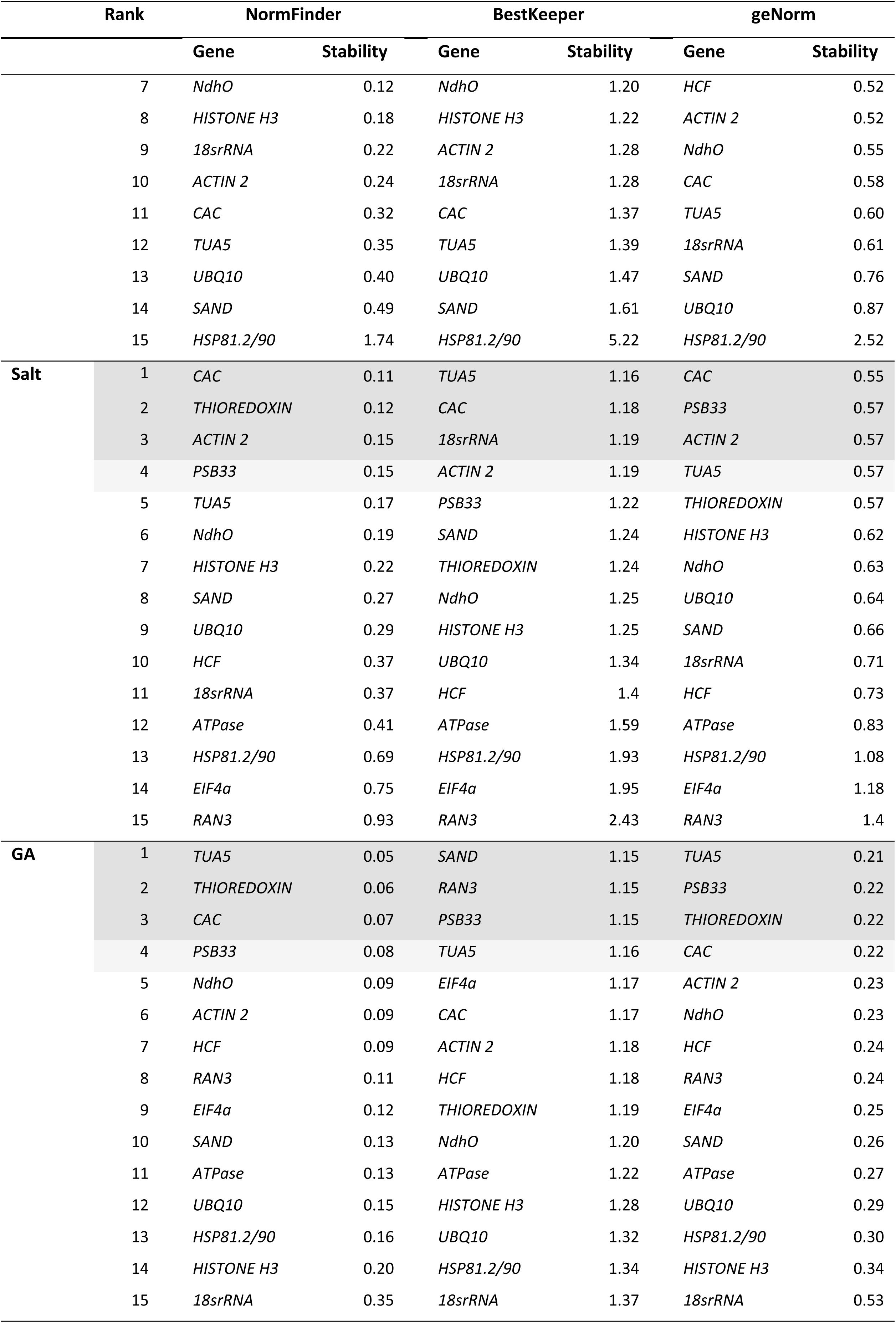
**Ranking of gene expression stability under stress conditions and hormone stimuli.** Genes were ranked using the three commonly used statistical algorithms NormFinder, BestKeeper and geNorm. The stability value describes the variance (NormFinder and geNorm) or standard deviation (BestKeeper).

**Table 4.**
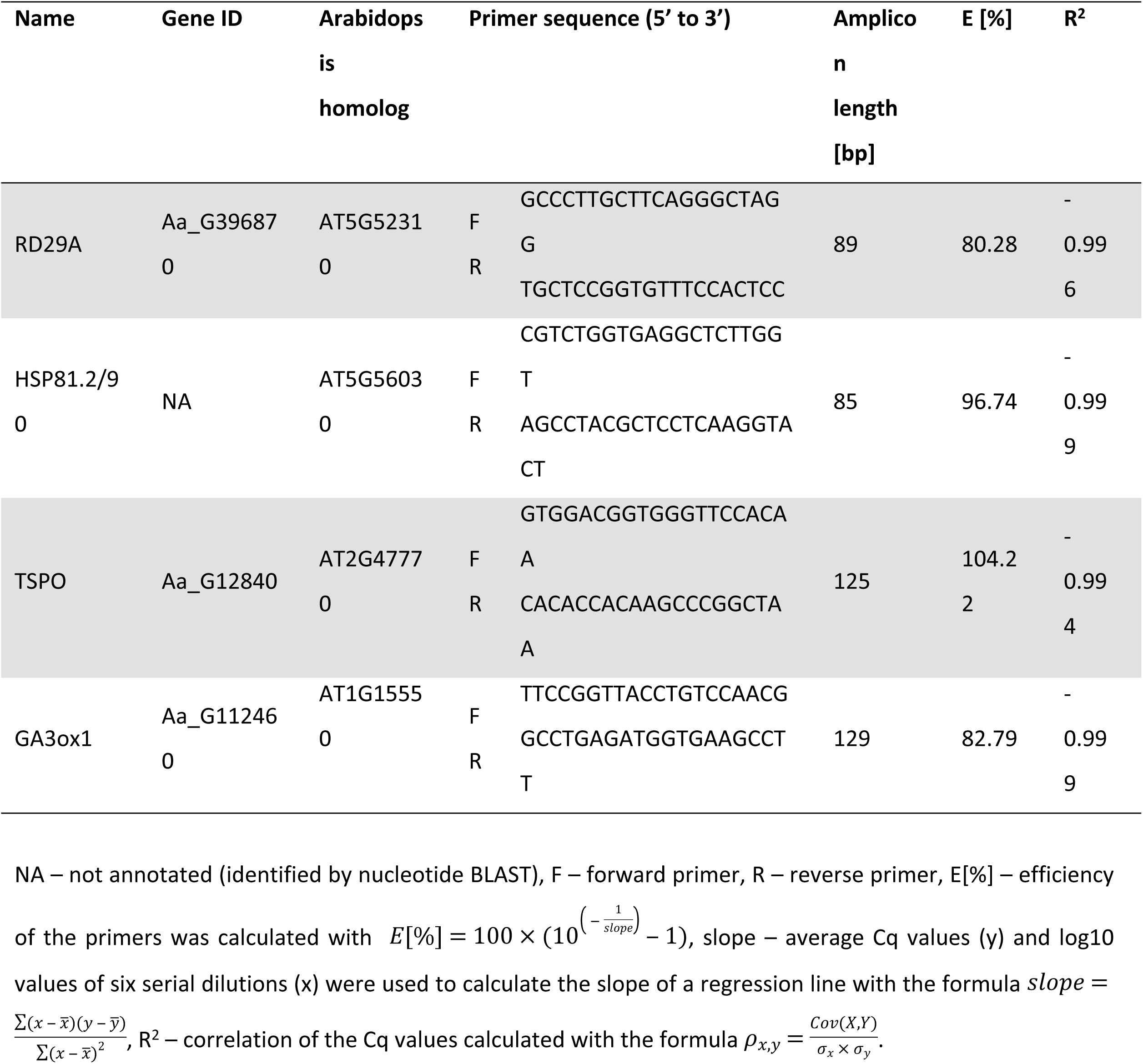
**Abiotic stress and hormone responsive genes, primers and amplicons.**

Normalisation of the results was carried out with one or two of the most stable reference genes determined in this study (Fig 3). For cold, normalisation was carried out with *RAN3* and/or *HCF*. Drought samples were normalised to *UBQ10* and/or *TUA5*. G*A* and salt responsive genes were normalised with *PSB33* and/or *TUA5.* Finally, the heat samples were normalised with *RAN3* and/or *PSB33.* The results clearly show that all treatments were successful, leading to an increased (cold, drought, heat, salt) or decreased (GA) expression of the responsive genes. Individual normalisation with each reference gene led to differences in the calculated fold changes of 28.9 % (cold), 7.5 % (drought), 3.6 % (GA), 9.3 % (heat) and 10.4 % (salt) between reference gene 1 and 2, respectively. These results clearly show the necessity to use two reference genes simultaneously.

**Fig 3.**
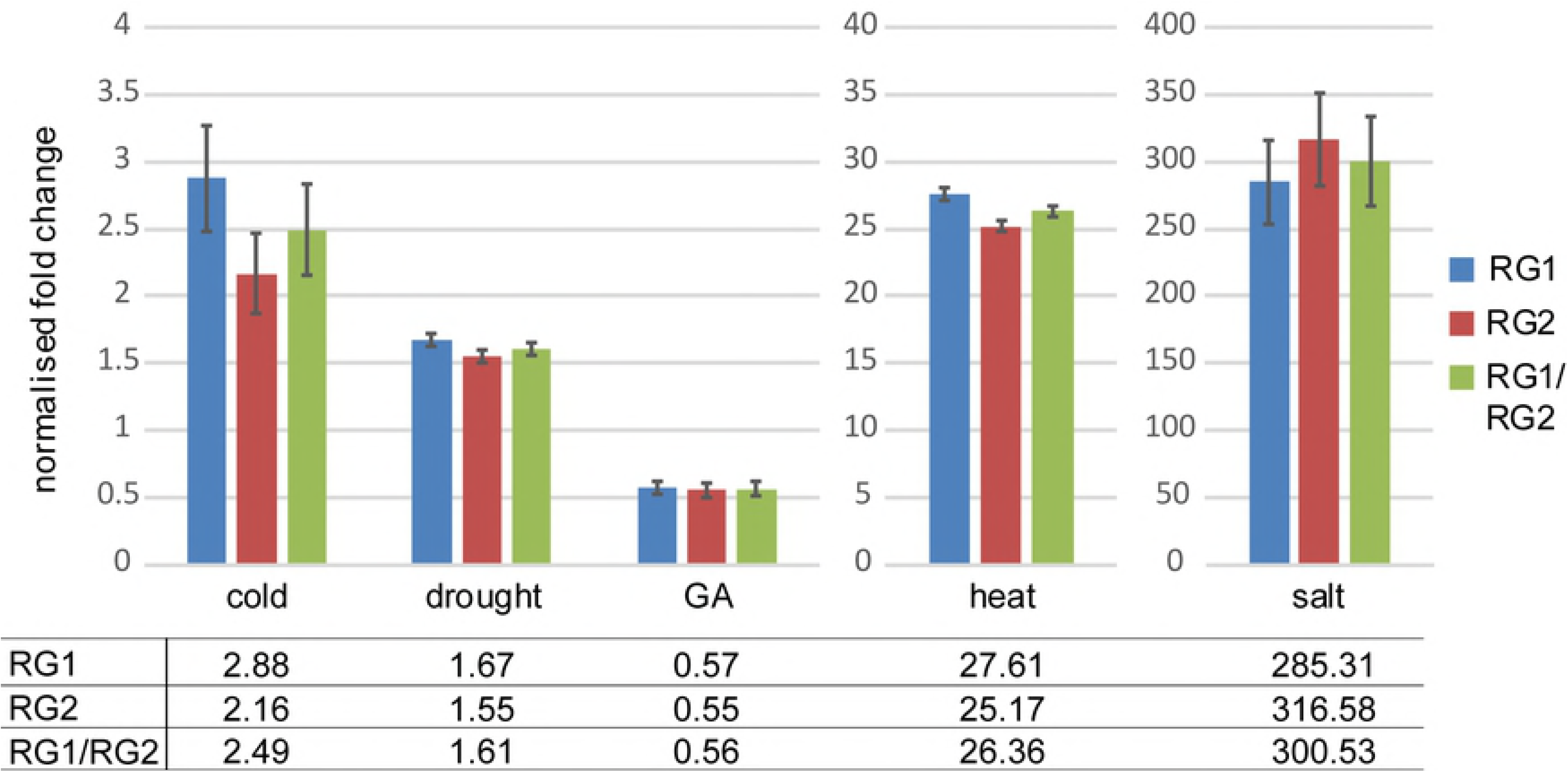
Comparison of specific stress response genes normalised with different reference genes. Normalisation of the stress response genes RD29A (RESPONSIVE TO DESICCATION 29A, cold and drought responsive), HSP81.2/90 (heat responsive), TSPO (OUTER MEMBRANE TRYPTOPHAN-RICH SENSORY PROTEIN-RELATED, salt responsive) and GA3ox1 (GIBBERELLIN 3-OXIDASE 1, GA responsive) was carried out with one or two reference genes (RG1 and RG2): cold – RAN3 and HCF, drought – UBQ10 and TUA5, GA and salt – PSB33 and TUA5, heat – RAN3 and PSB33.

## Material and Methods

### Plant growth conditions

For the abiotic stress treatments, seeds were surface sterilised with increasing concentrations of ethanol and grown on MS (10) plates without additional sucrose. The seeds were stratified for five days. Subsequently, plants were grown under long day conditions (16h light/8h darkness) at 21±1°C and 100±20 μmol/m^2^s light intensity.

The heat and cold treatments were carried out by transferring 5-7 day old seedlings to 38°C and 4°C, respectively. Samples were taken after 2 h. Drought was induced on MS plates which were exposed to liquid MS containing 20 % PEG 8000 for 24 h prior to the experiment. Samples were taken after 24 h. For salt treatments, two to three 5-7 day old seedlings were transferred from plates to liquid ½ MS (control) or liquid ½ MS containing 125 mM NaCl for 4 h under constant shaking.

For the GA treatment, plants were grown on soil under long day conditions (16h light/8h darkness) at 21±1°C and 200±20 μmol/m^2^s light intensity. The treatment was started right after germination and continued twice per week. Plants were sprayed with 20 μM GA4 (Sigma Aldrich, stock solution: 100 mM GA4 in EtOH, 0.1 % Silwet L-77 Loveland industries) or mock (0.1 % EtOH, 0.1 % Silwet). Leaves were harvested 14 days after germination at Zeitgeber time (ZT) 8.

All experiments were performed in three independent biological replicates. All samples were immediately frozen in liquid nitrogen and stored at -80°C.

### RNA extraction and cDNA synthesis

The frozen seedling samples were ruptured using a TissueLyser (Qiagen) and total RNA extraction was carried out with Tri-Reagent (Ambion by Life Technologies). All samples were treated with DNAseI (Thermo Fisher Scientific). The frozen leaf samples were ruptured using a TissueLyser (Qiagen), extracted with the RNAeasy Plant Mini Kit (Qiagen) and treated with DNAse (Ambion by Life Technologies). RNA integrity was controlled on a bleach gel (Aranda et al., 2012) and RNA concentration and purity was measured using a photometer (Eppendorf). cDNA was synthesised from 500 ng total RNA with oligodT primers, using the RevertAid First Strand cDNA Synthesis Kit (Thermo Fisher Scientific) according to the provided protocol. The cDNA was tested for DNA contamination and integrity by PCR and subsequent gel electrophoresis. The cDNA was diluted (1:10, 1:20, 1:40, 1:80, 1:160, and 1:320) to analyse primer efficiency and determine the correlation coefficient.

### Choice of candidate genes and primer design

The eight reference gene candidates *EIF4a, ACTIN 2, CAC, TUA5, HISTONE H3, HSP81.2/90, 18srRNA* and *SAND* were chosen due to their stable expression in other species (12–16). *PSB33, ATPase, THIOREDOXIN, HCF* and *NdhO* were chosen regarding their robust expression levels in Arabis in a time-course RNAseq experiment (data provided by Eva Willing, MPIPZ, Cologne) and/or due to their essential function for the plant. The *RAN3* primers were taken from Wang et al., 2009. *UBQ10* primers were kindly provided by Pan Pan Jiang.

Sequences were taken from TAIR (17), NCBI (National Centre for Biotechnology Information, www.ncbi.nlm.nih.gov) and the Genomic resources for *Arabis alpina* website (www.arabis-alpina.org; Willing et al., 2015). *In silico* sequence analysis was carried out with CLC DNA Workbench version 5.6.1. The primers were designed using GenScript Real-time PCR Primer Design (www.genscript.com) at an optimum Tm of 60±2°C. Amplicons showed a single band of the expected size in gel electrophoresis and a single peak in the melting curve (Fig S1). The PCR products were sequenced by GATC/ Eurofins Genomics to verify specific amplification.

### Quantitative real-time PCR (qPCR)

qPCRs were carried out in a QuantStudio 5 System (ABI/Life Technologies) equipped with the QuantStudio TM Design and Analysis Software version 1.4.1. The qPCRs were performed using plates (96 well, 0.2 ml) and cover foil (Opti-Seal Optical Disposable Adhesive) from BIOplastics. Reaction mixtures of 20 µl were composed from 5 µl SYBR Green (Thermo Fisher Scientific), 0.2 µl of each primer, 1 µl cDNA and 1 µl ddH_2_O. Amplification was carried out with the standard settings of the QuantStudio 5 System (50°C for 2 min, 95°C for 10 sec, 40 cycles at 95°C for 15 sec and 60°C for 1 min, followed by 95°C for 15 sec and a final dissociation curve from 60°C to 95°C). For each reference gene sample, three biological and three technical replicates were analysed. The impact of normalisation on stress/hormone responsive genes was analysed for three technical replicates of one biological replicate.

### Analysis of qPCR data

Efficiency calculations were carried out manually using Excel 2007. Extreme outliers were removed manually, a standard deviation of technical replicates was accepted below 0.5 Cq. Efficiency of primers was calculated in cDNA dilution series. The average Cq values (y) and log10 values of the dilutions (x) were used to calculate the slope of a regression line with the formula 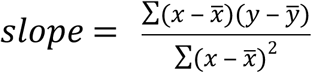. The slope was then used to calculate the efficiency of the primers 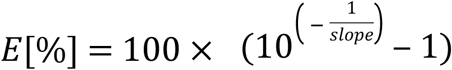. The correlation R^2^ of the values was calculated using the formula 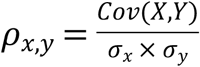. Primers for reference genes were accepted with an efficiency of 90-110 % and a correlation between -1 and -0.99. Primers for genes of interest were accepted with an efficiency of 80-120 % and a correlation between -1 and -0.99.

The stability of the reference genes within a given set of different treatments was calculated from the efficiency corrected data using the algorithms NormFinder (7), BestKeeper (8) and geNorm (9). The geNorm algorithm was also used to define the number of reference genes necessary for normalisation.

Normalisation against one reference gene was carried out using the normalisation factor, normalisation against two reference genes was carried out using the geometric mean of the normalisation factors, according to the geNorm manual (9). In accordance with this manual, standard deviations between biological replicates were calculated over the means of the single replicates, rather than the raw data.

## Discussion

Gene expression analysis by qPCR is a high-throughput method, which is considered to be very sensitive and reproducible. However, the accuracy of the results strongly depends on the experimental design, adequate normalisation and exact analysis of the produced data (18). Moreover, the qPCR primers must be specific, efficient, and - in the case of reference genes - stable in the tested conditions (18). In this study, we analysed primers for one established reference gene (*RAN3*; Wang et al., 2009) and 14 novel reference genes for *Arabis alpina* in several abiotic stress and hormone treatments. *EIF4a, ACTIN 2, CAC, TUA5, HISTONE H3, HSP81.2/90* and *SAND* were chosen because they were already established as reference genes in other species (12–16,19,20). In addition, we considered *PSB33, ATPase, THIOREDOXIN, HCF* and *NdhO*. The reference genes selected here are involved in various basic cellular functions including translation, proton transport, photosynthesis, protein degradation and cytoskeletal organisation. Our data suggest that all genes reported here are appropriate reference genes for the used tissues under non-stress conditions. As *PSB33* and *HCF* are functionally related to photosynthesis, they may be more appropriate for photosynthetic tissues.

The reference gene primers presented here have efficiency values between 96.42 and 107.01 %. Although these are very good efficiencies, it is essential to correct the qPCR results with these values because of the non-linearity of the PCR amplification steps (21). Consequently, we used efficiency-corrected data for identifying the most stable reference genes and for the analysis of the impact on normalisation. There is currently no consensus in the community, which of the three statistical algorithms – geNorm, NormFinder or BestKeeper – is most appropriate. One advantage of the geNorm algorithm is that it can be used for small sample sizes (22). However, the geNorm method is biased towards genes that are co-regulated (23). The algorithm takes into consideration, whether genes show a similar expression pattern (24), since it is assumed that similar changes in the expression of two independent genes reflect technical differences, such as the cDNA concentration, rather than changes caused by a treatment. By contrast, NormFinder considers variations across subgroups (7). The algorithm assumes that there is no systematic variation of the average of the tested samples, which can lead to a preference for reference genes with similar systematic variation (23). BestKeeper takes the standard deviation of each individual reference gene into account, which is an advantage over the other methods. The disadvantage of this algorithm is the use of a parametric method (Pearson correlation), which requires normally distributed data with a homogenous variance (8), which is not always the case. Additionally, BestKeeper uses the raw Cq values, while geNorm and Normfinder require normalised quantities. Therefore, the results obtained with BestKeeper are often different from those of the other two methods.

It is generally recommended to use more than one reference gene to guarantee optimal normalisation (18). Our data support this view. The normalised fold change values varied up to 28.9 % between two references. Moreover, we recommend that at least one of the reference gene amplicons contains introns at the genomic level to recognize potential contaminations with genomic DNA.

For cold treatments, we recommend *RAN3*, as the amplicon contains an intron, combined with *HCF* or *PSB33*. As *HCF* and *PSB33* are photosynthesis-associated proteins located in the thylakoid membrane, there is no obvious reason to prefer one over the other. For drought treatments, *UBQ10* and *TUA5* were the most stable transcripts, with *TUA5* also containing an intron. In heat, *RAN3* can be recommended in combination with *PSB33* or *EIF4a*. As setups for heat treatments often go along with specific light settings, it might be advisable to use *EIF4a* as a second reference here. For the salt treatment, we found that *CAC, TUA5* and *ACTIN 2* are suitable, intron containing reference genes, which can be combined with each other or *PSB33*. The combination *TUA5* and *PSB33* might be the most efficient choice, as it is also the best option for GA treatments.

We found that there is no single reference gene, which is the best choice for all treatments. However, all genes tested in this study, except for *HSP81.2/90* under heat conditions, meet the requirements necessary for adequate normalisation. Thus, this study provides data for the selection of suitable reference gene combinations for *Arabis alpina* in cold, drought, heat, salt and GA treatments. With this, we enable adequate normalisation in qPCR experiments under these conditions and provide novel reference genes for future experiments addressing other stresses or stimuli.

## Acknowledgements

We thank Pan Pan Jiang and Eva Willing for providing us with primer sequences and expression data, respectively. We also thank Jessica Pietsch and Tamara Gigolashvili for critically reading the manuscript.

## Supplementary Materials

**Table S1. Raw Cq values of treatments with cold, drought, heat, salt and gibberellic acid. Fig S1. Melting curves of candidate reference genes and stress/hormone responsive genes.**

**Fig S2. Standard curves of candidate reference genes and stress/hormone responsive genes.**

**Table S2. Ranking of gene expression stability under abiotic stress conditions and the combination of all treatments.** Genes were ranked using the three commonly used statistical algorithms NormFinder, BestKeeper and geNorm. The stability value describes the variance (NormFinder and geNorm) or standard deviation (BestKeeper).

## References

1. Wang R, Farrona S, Vincent C, Joecker A, Schoof H, Turck F, et al. PEP1 regulates perennial flowering in Arabis alpina. Nature [Internet]. 2009 May 15 [cited 2018 May 7];459(7245):423–2. 7. Available from: http://www.ncbi.nlm.nih.gov/pubmed/19369938

2. Willing E-M, Rawat V, Mandáková T, Maumus F, James GV, Nordström KJV, et al. Genome expansion of Arabis alpina linked with retrotransposition and reduced symmetric DNA methylation. Nat Plants [Internet]. Nature Publishing Group; 2015 Feb 2 [cited 2018 Dec 3];1(2):14023. Available from: http://www.nature.com/articles/nplants201423

3. Meusel H, JÄger EJ, Weinert E. Vergleichende Chorologie der Zentraleuropaeischen Flora. Jena, Germany: Fischer Verlag; 1965.

4. Koch MA, Kiefer C, Ehrich D, Vogel J, Brochmann C, Mummenhoff K. Three times out of Asia Minor: the phylogeography of Arabis alpina L. (Brassicaceae). Mol Ecol [Internet]. Wiley/Blackwell (10.1111); 2006 Feb 23 [cited 2018 Jul 30];15(3):825–39. Available from: http://doi.wiley.com/10.1111/j.1365-294X.2005.02848.x

5. Beilstein MA, Nagalingum NS, Clements MD, Manchester SR, Mathews S. Dated molecular phylogenies indicate a Miocene origin for Arabidopsis thaliana. Proc Natl Acad Sci U S A [Internet]. 2010 Oct 26 [cited 2017 Apr 30];107(43):18724–8. Available from: http://www.pnas.org/cgi/doi/10.1073/pnas.0909766107

6. Chopra D, Wolff H, Span J, Schellmann S, Coupland G, Albani MC, et al. Analysis of TTG1 function in Arabis alpina. BMC Plant Biol [Internet]. BioMed Central; 2014 [cited 2016 Aug 3];14(1):16. Available from: http://bmcplantbiol.biomedcentral.com/articles/10.1186/1471-2229-14-16

7. Andersen CL, Jensen JL, Ørntoft TF. Normalization of real-time quantitative reverse transcription-PCR data: a model-based variance estimation approach to identify genes suited for normalization, applied to bladder and colon cancer data sets. Cancer Res [Internet]. 2004 Aug 1 [cited 2018 May 6];64(15):5245–50. Available from: http://cancerres.aacrjournals.org/lookup/doi/10.1158/0008-5472.CAN-04-0496

8. Pfaffl MW, Tichopad A, Prgomet C, Neuvians TP. Determination of stable housekeeping genes, differentially regulated target genes and sample integrity: BestKeeper – Excel-based tool using pair-wise correlations. Biotechnol Lett [Internet]. Kluwer Academic Publishers; 2004 Mar [cited 2018 May 6];26(6):509–15. Available from: http://link.springer.com/10.1023/B:BILE.0000019559.84305.47

9. Vandesompele J, De Preter K, Pattyn F, Poppe B, Van Roy N, De Paepe A, et al. Accurate normalization of real-time quantitative RT-PCR data by geometric averaging of multiple internal control genes. Genome Biol [Internet]. BioMed Central; 2002 Jun 18 [cited 2018 May 6];3(7):research0034.1. Available from: http://genomebiology.biomedcentral.com/articles/10.1186/gb-2002-3-7-research0034

10. Murashige T, Skoog F. A Revised Medium for Rapid Growth and Bio Assays with Tobacco Tissue Cultures. Physiol Plant [Internet]. Wiley/Blackwell (10.1111); 1962 Jul 1 [cited 2018 May 6];15(3):473–97. Available from: http://doi.wiley.com/10.1111/j.1399-3054.1962.tb08052.x

11. Aranda PS, LaJoie DM, Jorcyk CL. Bleach gel: a simple agarose gel for analyzing RNA quality. Electrophoresis [Internet]. NIH Public Access; 2012 Jan [cited 2018 Aug 2];33(2):366–9. Available from: http://www.ncbi.nlm.nih.gov/pubmed/22222980

12. Tian C, Jiang Q, Wang F, Wang G-L, Xu Z-S, Xiong A-S. Selection of Suitable Reference Genes for qPCR Normalization under Abiotic Stresses and Hormone Stimuli in Carrot Leaves. Jain M, editor. PLoS One [Internet]. Public Library of Science; 2015 Feb 6 [cited 2017 Mar 2];10(2):e0117569. Available from: http://dx.plos.org/10.1371/journal.pone.0117569

13. Wang H, Wang J, Jiang J, Chen S, Guan Z, Liao Y, et al. Reference genes for normalizing transcription in diploid and tetraploid Arabidopsis. Sci Rep [Internet]. Nature Publishing Group; 2015 May 27 [cited 2018 Aug 2];4(1):6781. Available from: http://www.nature.com/articles/srep06781

14. Demidenko N V., Logacheva MD, Penin AA. Selection and Validation of Reference Genes for Quantitative Real-Time PCR in Buckwheat (Fagopyrum esculentum) Based on Transcriptome Sequence Data. Mariño-Ramírez L, editor. PLoS One [Internet]. Public Library of Science; 2011 May 12 [cited 2018 Aug 2];6(5):e19434. Available from: http://dx.plos.org/10.1371/journal.pone.0019434

15. Czechowski T, Stitt M, Altmann T, Udvardi MK, Scheible W-R. Genome-wide identification and testing of superior reference genes for transcript normalization in Arabidopsis. Plant Physiol [Internet]. American Society of Plant Biologists; 2005 Sep [cited 2017 Jun 9];139(1):5–17. Available from: http://www.ncbi.nlm.nih.gov/pubmed/16166256

16. Wan Q, Chen S, Shan Z, Yang Z, Chen L, Zhang C, et al. Stability evaluation of reference genes for gene expression analysis by RT-qPCR in soybean under different conditions. Li X, editor. PLoS One [Internet]. 2017 Dec 13 [cited 2018 Nov 28];12(12):e0189405. Available from: http://www.ncbi.nlm.nih.gov/pubmed/29236756

17. Berardini TZ, Reiser L, Li D, Mezheritsky Y, Muller R, Strait E, et al. The arabidopsis information resource: Making and mining the “gold standard” annotated reference plant genome. genesis [Internet]. 2015 Aug 1 [cited 2018 Jan 25];53(8):474–85. Available from: http://doi.wiley.com/10.1002/dvg.22877

18. Bustin SA, Benes V, Garson JA, Hellemans J, Huggett J, Kubista M, et al. The MIQE Guidelines: Minimum Information for Publication of Quantitative Real-Time PCR Experiments. Clin Chem [Internet]. 2009 [cited 2017 Aug 29];55(4). Available from: http://clinchem.aaccjnls.org/content/55/4/611/tab-figures-data

19. Wu J, Zhang H, Liu L, Li W, Wei Y, Shi S. Validation of Reference Genes for RT-qPCR Studies of Gene Expression in Preharvest and Postharvest Longan Fruits under Different Experimental Conditions. Front Plant Sci [Internet]. 2016 Jun 3 [cited 2018 Nov 28];7:780. Available from: http://www.ncbi.nlm.nih.gov/pubmed/27375640

20. Najafpanah MJ, Sadeghi M, Bakhtiarizadeh MR. Reference Genes Selection for Quantitative Real-Time PCR Using RankAggreg Method in Different Tissues of Capra hircus. Niemann H, editor. PLoS One [Internet]. 2013 Dec 16 [cited 2018 Nov 28];8(12):e83041. Available from: http://www.ncbi.nlm.nih.gov/pubmed/24358246

21. Ruijter JM, Ramakers C, Hoogaars WMH, Karlen Y, Bakker O, van den Hoff MJB, et al. Amplification efficiency: linking baseline and bias in the analysis of quantitative PCR data. Nucleic Acids Res [Internet]. 2009 Apr 1 [cited 2018 Dec 13];37(6):e45–e45. Available from: http://www.ncbi.nlm.nih.gov/pubmed/19237396

22. Serrano M, Moreno-Sánchez N, González C, Marcos-Carcavilla A, Van Poucke M, Calvo JH, et al. Use of Maximum Likelihood-Mixed Models to select stable reference genes: a case of heat stress response in sheep. BMC Mol Biol [Internet]. 2011 Aug 17 [cited 2018 Dec 9];12(1):36. Available from: http://bmcmolbiol.biomedcentral.com/articles/10.1186/1471-2199-12-36

23. Mehdi Khanlou K, Van Bockstaele E. A critique of widely used normalization software tools and an alternative method to identify reliable reference genes in red clover (Trifolium pratense L.). Planta [Internet]. 2012 Nov 21 [cited 2018 Dec 9];236(5):1381–93. Available from: http://www.ncbi.nlm.nih.gov/pubmed/22718310

24. Vandesompele J, De Preter K, Pattyn F, Poppe B, Van Roy N, De Paepe A, et al. Accurate normalization of real-time quantitative RT-PCR data by geometric averaging of multiple internal control genes. Genome Biol [Internet]. BioMed Central; 2002 Jun 18 [cited 2018 Apr 19];3(7):RESEARCH0034. Available from: http://www.ncbi.nlm.nih.gov/pubmed/12184808

